# Hidden infections: Linking parasite ecology, fish health and stakeholder perceptions in Nile tilapia aquaculture in central Kenya

**DOI:** 10.64898/2026.11.04.764176

**Authors:** Miriam Isoyi Shigoley, Nicolas Antoine-Moussiaux, Thierry Jauniaux, Thomas-Julian Irabor, Gyrhaiss Kapepula Kasembele, Daniel Mungai Ndegwa, Bram J. Vanschoenwinkel, Maarten P. M. Vanhove

**Affiliations:** Research Group Zoology: Biodiversity and Toxicology, Centre for Environmental Sciences, Hasselt University, 3590 Diepenbeek, Belgium; Department of Veterinary Management of Animal Resources, Faculty of Veterinary Medicine, University of Liège, 4000 Liège, Belgium; Department of Public Health, Faculty of Medicine, University of Liège, 4000 Liège, Belgium; Department of Morphology and Pathology, Faculty of Veterinary Medicine, University of Liège, 4000 Liège, Belgium; Unité de Recherche en Biodiversité et Exploitation durable des Zones Humides (BEZHU), Faculté des Sciences Agronomiques, Université de Lubumbashi, Haut-Katanga, 1825, Democratic Republic of Congo; Kenya Fisheries Service, National Aquaculture Technology Development and Innovations Transfer Centre, 26-10230 Sagana, Kenya; Department of Biology, Vrije Universiteit Brussel (VUB), 1050, Brussels, Belgium; Centre for Environmental Management, University of the Free State, 339-9300, Bloemfontein, South Africa; Royal Belgian Institute of Natural Sciences, OD Natural Environment, Freshwater Biology, 1000 Brussels, Belgium

**Keywords:** Fish parasitology, Diagnostics, Host-parasite interactions, Aquaculture, Histopathology

## Abstract

Parasite infections in farmed fish may remain unrecognised when fish-health assessment relies mainly on visible signs. We investigated metazoan parasite occurrence, parasite diversity and community composition, associations between gill-parasite burden and histological alterations, and stakeholder recognition of fish-health problems in Nile tilapia (*Oreochromis niloticus*) aquaculture in central Kenya. We examined 157 fish from 20 farms, assessed gill histology in 137 fish, measured water quality once per farm, and collected questionnaire and interview data from 48 stakeholders. Five metazoan parasite groups were recorded, with monopisthocotylans the most prevalent (36.3%). Higher monopisthocotylan burden was associated with a greater cumulative number of gill lesion types (odds ratio 1.86, 95% CI 1.43–2.42, p < 0.001), whereas copepod burden was not. Parasite diversity was higher in the larger operational pond category across all three Hill-number orders (p = 0.006–0.008), and parasite-community composition differed between categories (PERMANOVA, p = 0.034), although these comparisons were exploratory because only four farms were in the larger category and farm category overlapped with county. Single-time-point water-quality measurements showed no clear associations with parasite burden, diversity or community composition after multiple-testing correction. Farmers and fisheries or extension officers recognised signs of poor fish health but rarely identified parasites as a specific cause, and routine diagnostic confirmation was uncommon. The findings reveal a gap between laboratory detection of parasite-associated gill alterations and farm-level recognition and diagnosis.

## 1. Introduction

Aquaculture has expanded rapidly in response to growing demand for aquatic foods, increasing the importance of effective fish-health management. In 2024, global fisheries and aquaculture production reached 235 million tonnes, including 103 million tonnes of farmed aquatic animals (FAO, 2026). As production expands, disease recognition, prevention and access to diagnostic support become increasingly important, particularly where movement of fish seed and limited aquatic animal-health services can increase health risks (Bondad-Reantaso et al., 2005; Cameron, 2002).

Aquaculture development remains uneven across regions. In Africa, institutional and technical constraints, including limitations in fish-health management, continue to restrict development in many production systems. Kenya has nevertheless experienced substantial aquaculture expansion over the past two decades. Government investment between 2009 and 2013 supported pond construction, hatchery capacity, fingerlings, feeds and training for farmers and extension officers, contributing to the expansion of pond aquaculture (Munguti et al., 2021; Musa et al., 2012). Nile tilapia (*Oreochromis niloticus*) is among the most widely farmed fish and is important for food security and income generation, particularly in low- and middle-income countries (Meyer and Emam, 2024; WorldFish, 2022).

Fish parasites are an important but often overlooked aquaculture health challenge. Farmed tilapia can host monopisthocotylans, digeneans, cestodes, nematodes, acanthocephalans and parasitic crustaceans (Shinn et al., 2023). Some infections can affect growth, feeding efficiency and survival, particularly at high burdens or when fish experience additional stressors, while visible lesions can reduce market value and some fish-borne parasites have zoonotic importance (Assane et al., 2022; Chai et al., 2005; Dong et al., 2015; Wanja et al., 2020).

Parasite transmission and infection patterns can be influenced by environmental conditions and farm management. Water temperature, dissolved oxygen, conductivity and nutrient concentrations can affect parasite development, survival of free-living stages and host susceptibility, while stocking, pond design, fingerling source and biosecurity may alter exposure and transmission (Allah et al., 2021; Shinn et al., 2023). Connections between farms and natural water bodies can also facilitate exchange between cultured and wild fish populations (Bouwmeester et al., 2021; Johansen et al., 2011).

Evidence from Kenyan inland waters indicates that farmed and wild Nile tilapia harbour diverse parasite groups, including monopisthocotylans, digeneans, nematodes, acanthocephalans and parasitic crustaceans, some of which are associated with reduced fish health and aquaculture productivity (Kibet et al., 2019). In central Kenya, black spot infections caused by metacercariae of *Neascus* spp. have been reported together with tissue alterations and concurrent bacterial infections, including those of *Aeromonas* sp., *Klebsiella* sp. and *Micrococcus* sp. (Wanja et al., 2020), while clinostomid trematodes collected during the present sampling campaign were subsequently characterised morphologically and molecularly, including species of *Clinostomum* and *Euclinostomum* (Shigoley et al., 2025).

In this study, the term parasites refers to metazoan parasites, including helminths and parasitic crustaceans. These organisms are often difficult to detect without targeted examination and may remain unnoticed for extended periods because they are not always visible externally or associated with clear clinical signs (Bondad-Reantaso et al., 2005). Moreover, parasite presence does not necessarily indicate parasitic disease. Infections may remain subclinical or produce non-specific signs, and their health relevance depends on whether they are associated with tissue damage, impaired function or other measurable host effects (Bernet et al., 1999; Steckert et al., 2018). Parasites may therefore be overlooked during routine fish health assessment and receive limited attention in discussions of disease prevention and farm management (Kyule-Muendo et al., 2022; Shigoley et al., 2024). At farm level, recognition is further shaped by access to aquatic veterinary services, diagnostic laboratories and fish-health training. Farmers may therefore rely mainly on visual inspection and experience when responding to suspected disease, while extension officers may have limited specialised training in aquatic animal health (Dimelu et al., 2018; Kyule-Muendo et al., 2022; Ssekyanzi et al., 2023).

Parasite screening, gill histology and stakeholder accounts capture different aspects of fish health. Examining these components within the same study setting allows laboratory-detected infections and tissue alterations to be compared with what farmers and fisheries or extension officers recognise and act upon, without assuming a causal relationship between the biological and stakeholder findings.

This study, therefore, examined parasite occurrence, diversity, community composition, and associated gill alterations together with stakeholder recognition and management of fish-health problems. We hypothesised that: (1) parasite diversity and community composition would differ between operational farm categories; (2) higher gill-parasite burden would be associated with a greater number of gill lesion types; and (3) water-quality measurements taken at sampling would be associated with parasite burden, diversity or community composition. Because only four farms were classified as larger-scale and farm categories overlapped with county, the first hypothesis was interpreted as an exploratory comparison rather than a causal test of production scale. In parallel, the stakeholder component examined how farmers and fisheries or extension officers recognised and responded to fish-health problems.

## 2. Materials and methods

### 2.1. Study design and integration

Biological and stakeholder data were collected during the same field campaign and analysed separately. The findings were integrated during interpretation by comparing laboratory-detected parasites and gill alterations with stakeholder recognition, diagnostic practices and responses to fish-health problems.

### 2.2. Study area

The study was conducted between January and February 2024 at 20 Nile tilapia farms in Kiambu, Nyeri, Kirinyaga and Embu counties in central Kenya (Fig. 1). Aquaculture in the region is dominated by smallholder pond systems, although farms vary in pond number, production infrastructure and management intensity (Emam et al., 2024). Farms were identified with the assistance of county fisheries extension officers, whose local knowledge supported the selection of farms spanning this variation in production and management practices.

**Fig. 1.**
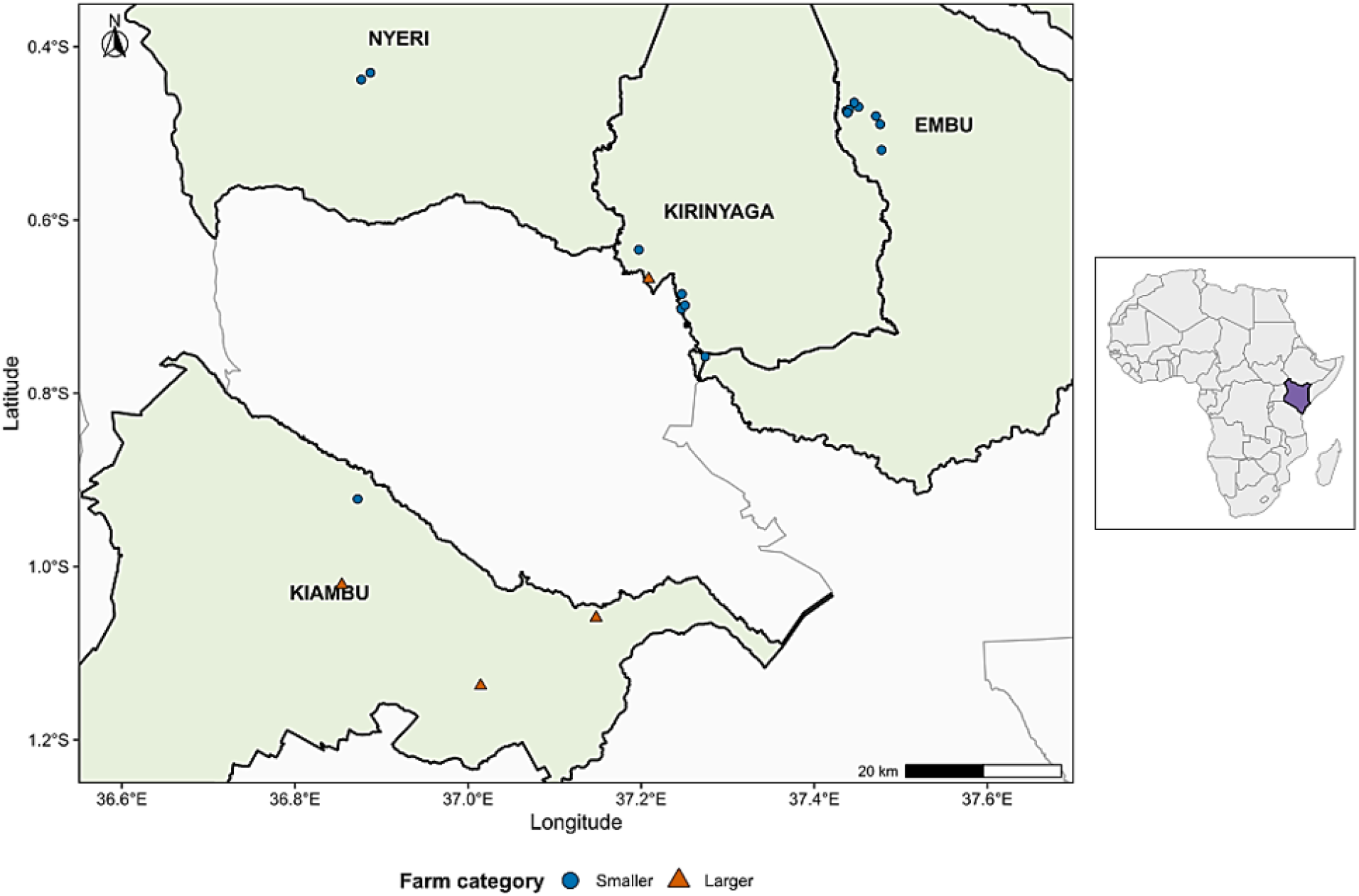
Location of the 20 Nile tilapia farms sampled in central Kenya. Study farms were located in Kiambu, Nyeri, Kirinyaga and Embu counties; blue circles indicate smaller farms and orange triangles larger farms. The map was produced in R version 4.5.2 (R Core Team, 2025) using RStudio version 2025.09.2+418 (Posit Software, Boston, MA, USA)

For analysis, farms were grouped according to the number of operational ponds. Farms with 1-3 operational ponds (n = 16) were classified as smaller-scale, whereas farms with more than three operational ponds were classified as larger-scale. Three of the larger-scale farms had 6,7 and 9 ponds while the fourth was an institutional farm reported to have between 100-150 ponds in total (Table 1). The terms smaller-scale and larger-scale are used only as labels for these two pond-number groups; they do not correspond to Kenyan or FAO definitions of farm scale, and pond number was not considered a measure of production intensity.

**Table 1.**
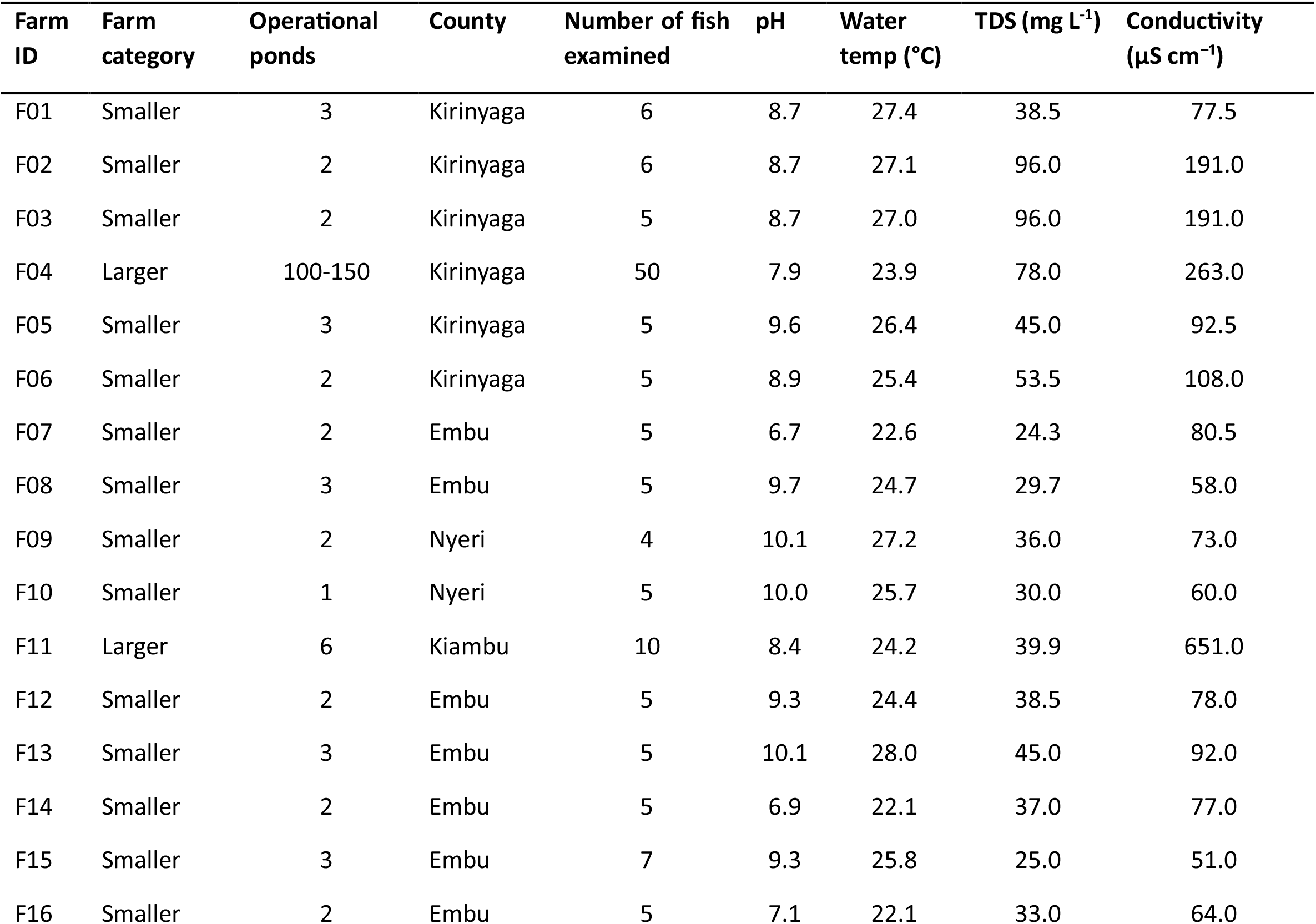
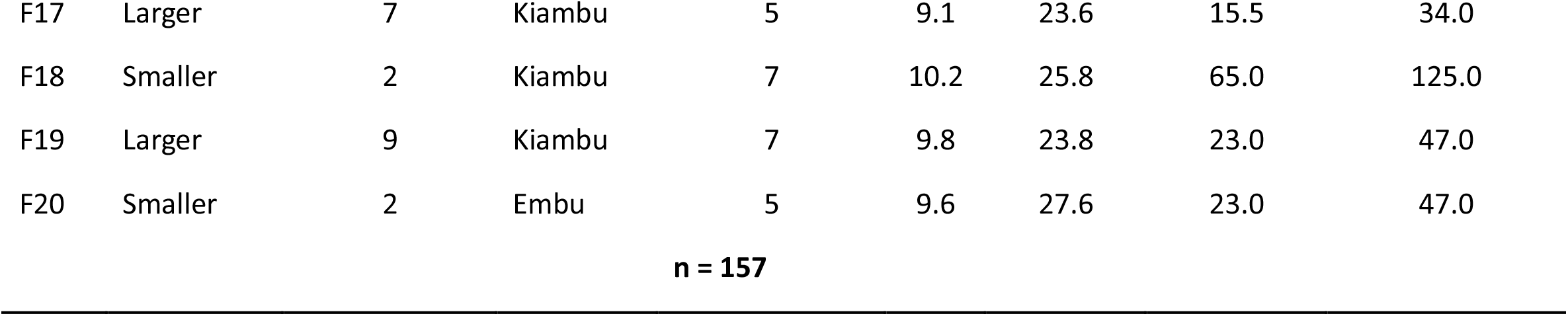
Number of fish examined per farm, operational farm category, county and water-quality measurements recorded at sampling.

The two groups were unevenly distributed across counties: three of the four larger-scale farms were in Kiambu, and one was in Kirinyaga, whereas all sampled farms in Nyeri and Embu were smaller-scale. County and farm categories were therefore partly confounded. Most sampled farms had reported total areas below 1,500 m^2^ and were consistent with smallholder pond systems described for Kenya (Coche et al., 1994). The institutional farm was the exception. A large number of specimens was collected at the institutional farm where more fish were available at the time of sampling. The field sampling was conducted after research clearance from the National Commission for Science, Technology and Innovation (NACOSTI); Permit No. NACOSTI/P/23/31261.

### 2.3. Water quality measurements

At the time of sampling, water temperature (°C), pH, conductivity (µS cm^−1^), and total dissolved solids (TDS; mg L^−1^) were measured in situ at each farm using a portable 4-in-1 water quality meter (Model EZ-9908; ESMART, China). The multi-meter was calibrated prior to deployment using commercial standard buffers (pH 4.01, 6.86, and 9.18) and a 1413 µS cm^−1^ conductivity reference solution. Each value is a single measurement per farm and does not represent seasonal or long-term conditions. These measurements were analysed as exploratory environmental correlates of parasite burden, diversity and community composition, and were not used as predictors of gill alterations.

### 2.4. Fish collection and parasite screening

Fish were collected from the farms using seine nets and euthanised by cervical dislocation. The intended number of fish per farm was ten, but the number obtained varied according to fish availability and farmer permission. Each fish was dissected, and the gills and intestinal tract were removed and preserved separately in 70% ethanol for subsequent parasitological examination. For gill-infecting parasites, the right gill arches were screened under a Leica EZ4 stereomicroscope (Leica Microsystems). Parasites were carefully removed using fine entomological needles and mounted on glass slides.

A small fin clip was taken from each individual and preserved in absolute ethanol for genetic reference. These samples were later deposited in the ichthyology collection of the Royal Belgian Institute of Natural Sciences, Belgium (AB49103238–285, AB42579285–332, AB42579752–764, AB42610341–388).

### 2.5. Parasite identification and molecular analyses

Recovered parasites were identified to the lowest possible taxonomic level using an integrative taxonomic approach involving group-appropriate morphological and molecular methods. Depending on the parasite group, these methods included light microscopy, molecular barcoding, and scanning electron microscopy (SEM). Whole parasite mount preparations were examined under an Olympus BX53F2 microscope (Olympus Soft Imaging Solutions GmbH, Johann-Krane-Weg, Münster, Germany), fitted with an Olympus DP23 camera.

Monopisthocotylans were fixed with Hoyer’s medium (Humason, 1979), covered with a coverslip and sealed after 24 hours with Glyceel (Bates, 1997). Their identification was based on the morphology of the sclerotized parts of the haptor and copulatory organs following Ergens (1981); Muterezi Bukinga et al. (2012); Pariselle et al. (2003); and Pariselle and Euzet (2009).

The digenean metacercariae collected during this sampling campaign were previously characterised using morphological and molecular approaches (Shigoley et al., 2025). Molecular identification was attempted for acanthocephalans but did not yield usable products. Nematodes were examined morphologically and assigned to Anisakidae using the identification keys and descriptions provided by (Moravec, 2019).

For copepods, molecular analyses targeted partial 18S and 28S rDNA and mitochondrial cytochrome *c* oxidase subunit I (COI mtDNA). Copepod DNA was extracted from egg strings using a protocol adapted from Kmentová et al. (2021). The 18S and 28S rDNA regions were amplified using primers described by Song et al. (2007), and COI was amplified using LCO1490 and HCO2198 (Folmer et al., 1994) primers. Amplification success was checked by gel electrophoresis, PCR products were purified using a GeneJet Purification Kit (Thermo Fisher Scientific, Waltham, MA, USA), and Sanger sequencing was performed by Macrogen (Amsterdam, The Netherlands). Detailed extraction and PCR conditions are provided in the Supplementary Methods. The sequences generated were compared with GenBank records using Basic Local Alignment Search Tool (BLAST; https://blast.ncbi.nlm.nih.gov/Blast.cgi) (Altschul et al., 1990). Reference sequences from closely related *Lamproglena* species were retrieved from GenBank, and alignments were generated using MUSCLE v5 under default parameters. Maximum-likelihood model selection and pairwise genetic-distance analyses were performed in MEGA v11.0.13. No usable 18S rDNA sequences were obtained; subsequent analyses therefore used 28S rDNA and COI mtDNA. Pairwise distances used K2P+G for 28S (Kimura, 1980) and GTR+G for COI. Sequences generated in this study were submitted to GenBank under accession numbers PZ790842–PZ790843 and PZ786304–PZ786311.

The mounted parasite specimens were deposited in the collection of the Research Group Zoology: Biodiversity and Toxicology at Hasselt University, Diepenbeek, Belgium (HU) – XXXVIII.3.28 - XLI.4.38.

### 2.6. Infection parameters and diversity indices

Infection parameters were calculated as described by Bush et al. (1997). For diversity analyses, parasites were grouped into 12 taxonomic units at the lowest identification level consistently available across the full dataset: eight monopisthocotylan species, *Lamproglena* sp., Digenea, *Acanthogyrus* cf. *tilapiae* and Anisakidae. Digenean metacercariae were treated as a single taxonomic unit because species-level identification was available only for a subset of specimens; detailed morphological and molecular identification is reported in Shigoley et al. (2025). Diversity was expressed as Hill numbers, reported as effective numbers of taxa: q = 0 represented taxon richness, q = 1 the exponential of Shannon diversity and q = 2 the inverse Simpson measure, 1/Σp_i_^2^ (Jost, 2006). Farms in which no parasites were detected were retained and assigned a value of zero at all three orders.

Because the number of fish examined differed among farms and was particularly high at F04, incidence-based rarefaction was used as a sampling-effort check for q = 0. Individual fish were treated as sampling units, and farm categories were compared at a common number of fish. Rarefaction was restricted to interpolation within the observed sampling range and was used only to assess whether differences in observed richness persisted after standardising sampling effort, following Gotelli and Colwell (2001).

### 2.7. Histopathological analysis

Separate gill tissue samples were fixed in buffered formalin, dehydrated through graded ethanol, cleared in xylene, embedded in paraffin wax, sectioned, mounted on glass slides and stained with haematoxylin and eosin. Gill sections were examined for five lesion types: epithelial hypertrophy, epithelial hyperplasia, lamellar fusion, vascular congestion of the secondary lamellae and inflammation. Epithelial hypertrophy was identified as enlargement of individual epithelial cells; epithelial hyperplasia as an increase in epithelial cell number resulting in thickening of the lamellar or interlamellar epithelium; lamellar fusion as partial or complete joining of adjacent secondary lamellae; vascular congestion as engorgement of lamellar vascular spaces with erythrocytes; and inflammation as inflammatory-cell infiltration within the gill tissue (Bernet et al., 1999; Wolf et al., 2015). These lesions have been reported in health assessments of farmed Nile tilapia but are not specific to parasite infection (Bernet et al., 1999; Steckert et al., 2018). Each lesion was recorded as present or absent. The number of lesion types present in each fish was summed to produce a cumulative count from 0 to 5; this measure represents the number of alteration types present and not their individual severity.

### 2.8. Stakeholder survey and interviews

Before the main fieldwork, a pilot study was conducted to refine the questionnaires and interview guides. An online survey created in Google Forms was distributed through a QR code at a scientific conference organised by the Pan African Fish and Fisheries Association (PAFFA) in 2023 and was voluntarily completed by 71 respondents working in aquaculture, fisheries, research, regulation, extension, industry, and environmental management. Five pilot interviews identified questions requiring clarification. Pilot survey and interview data were used only to revise the study instruments and were excluded from the main analysis.

During the main fieldwork, structured questionnaires were administered through KoboToolbox to 36 fish farmers, 11 fisheries or extension officers, and one veterinary officer. Semi-structured interviews examined recognition of fish-health problems, responses to suspected disease, information sources, access to diagnostic services and institutional support (Supplementary Interview Guide S1). Interviews were conducted in English or Kiswahili according to participant preference. Field notes taken during and immediately after each interview were organised and reviewed to document responses and relevant observations. Open-ended survey responses and interview field notes were analysed using a structured thematic approach, with initial codes grouped into broader themes concerning disease recognition, diagnostic practices, treatment decisions, institutional support and information flow (Braun and Clarke, 2006). Because the qualitative material consisted mainly of field notes rather than verbatim transcripts, it was used to describe recurring patterns and contextualise the biological findings, not to claim thematic saturation or conceptually interpret stakeholders’ views and logic. The single veterinary officer contributed contextual information and was not treated as a separate stakeholder group for comparative analysis.

### 2.9. Statistical analyses

All statistical analyses were performed in R version 4.5.2 (R Core Team, 2025) using RStudio 2025.09.2+418 (Posit Software, Boston, MA, USA). Community analyses were conducted using *vegan* (Oksanen et al., 2022), incidence-based rarefaction using *iNEXT* (Hsieh et al., 2016), and ordinal regression using *ordinal* (Christensen, 2026). Figures were produced using *ggplot2* (Wickham, 2016).

Given that only 20 farms were sampled and four were classified as larger-scale, farm-level analyses were restricted to the stated study questions and unnecessary model complexity was avoided. Farm-level Hill numbers were compared between operational farm categories using Wilcoxon rank-sum tests. The comparison was repeated after excluding F04 to assess whether the larger sample from that farm drove the result. Incidence-based rarefaction of q = 0 was calculated across individual fish for the full dataset and again after excluding F04. Parasite-community composition was analysed among parasite-positive farms using a square-root-transformed farm-by-taxon abundance matrix and Bray–Curtis dissimilarities. Principal coordinates analysis (PCoA) was used for visualisation. Differences between farm categories were tested using PERMANOVA with 9,999 permutations, and homogeneity of multivariate dispersion was checked with PERMDISP. Community analyses were repeated after excluding F04 and F11 separately.

Water-quality associations were examined at farm level. Spearman correlations tested each water variable separately against mean parasite abundance per fish and each Hill number. Separate one-variable PERMANOVAs tested associations with community composition. P-values were adjusted using the Benjamini–Hochberg false-discovery-rate procedure separately for tests of parasite burden, Hill diversity and community composition. These analyses used the single measurements taken at sampling and were interpreted only as associations with conditions at that time. Water-quality measurements were not used as predictors of gill alterations because only one measurement was available per farm and could not represent the longer-term exposure over which tissue alterations may have developed.

The cumulative lesion-type count was analysed as an ordered outcome using a proportional-odds ordinal logistic model. Predictors were ln(monopisthocotylan abundance + 1), ln(copepod abundance + 1), host sex, operational farm category and standardised total length. The proportional-odds assumption was checked using a nominal-effects test. To assess clustering of fish within farms, the model was repeated with farm identity included as a fixed effect; farm category was omitted from this sensitivity model because it did not vary within farms. A linear model with farm-clustered standard errors was used only as a secondary sensitivity check. Each individual lesion type was analysed separately using binary logistic regression with the same parasite and host predictors; false-discovery-rate adjustment was applied across the five lesion tests for each parasite group. These lesion-specific models were treated as secondary analyses and are reported in the Table S2.

### 2.10. Ethical considerations

The stakeholder study was reviewed by the Hasselt University Social and Societal Ethics Committee (SMEC), which granted an exemption from formal ethical approval. Participants were informed about the study purpose, voluntary participation, their right to decline questions or withdraw, and the intended use of responses and photographs. Verbal informed consent was obtained before questionnaires, interviews and photographs. Personal identifiers were removed from the analytical dataset, access to the data was restricted to the research team, and the data were stored and retained in accordance with the institution’s data-management procedures and the General Data Protection Regulation.

## 3 Results

### 3.1. Study overview and water conditions

A total of 157 Nile tilapia were examined for ecto- and endoparasites across 20 farms (Table 1). The number of fish examined ranged from four to ten at most farms, while 50 fish were examined at F04 because more fish were available at this government-operated facility. The stakeholder component comprised 48 participants: 36 farmers, 11 fisheries or extension officers and one veterinary officer. Most participating farmers (72%) owned the farms they managed, and their reported aquaculture experience ranged from one to 24 years.

At the time of sampling, mean pH was 8.94 (range 6.7–10.2), mean water temperature was 25.2 °C (22.1–28.0 °C), mean TDS was 43.6 mg L^−1^ (15.5–96.0 mg L^−1^), and mean conductivity was 123 µS cm^−1^ (34.0–651 µS cm^−1^). These values describe the measurements taken at the time of sampling and not seasonal or long-term conditions.

### 3.2. Parasite occurrence

Five metazoan parasite groups were recorded (Table 2). Monopisthocotylans were the most prevalent, infecting 57 of 157 fish (36.3%), followed by copepods (34/157; 21.7%), digeneans (27/157; 17.2%), acanthocephalans (10/157; 6.4%) and nematodes (2/157; 1.3%). At farm level, monopisthocotylans occurred in 14/20 farms (10 smaller-scale and all four larger-scale farms), copepods in 3/20 farms, digeneans in 2/20 farms, acanthocephalans in 2/20 farms and nematodes in 1/20 farms. Copepods, digeneans, acanthocephalans and nematodes were detected only in farms in the larger-scale category.

**Table 2.** Infection parameters for the five parasite groups recorded in *Oreochromis niloticus*.

| **Parasite group** | **Infected hosts** | **Positive farms** | **Prevalence (P) % (95% CI)** | **Mean intensity (MI) (range)** |
|---|---|---|---|---|
| Monopisthocotylans | 57/157 | 14/20 | 36.3 (28.8–44.3) | 39.9 (1-930) |
| Copepods | 34/157 | 3/20 | 21.7 (15.5–28.9) | 4.7 (1-21) |
| Digeneans | 27/157 | 2/20 | 17.2 (11.6–24.0) | 7.3 (1-38) |
| Acanthocephalans | 10/157 | 2/20 | 6.4 (3.1–11.4) | 1.6 (1-4) |
| Nematodes | 2/157 | 1/20 | 1.3 (0.2–4.5) | 1.0 (1-1) |

A total of 2,275 monopisthocotylans were recovered from the gills, representing eight species of *Cichlidogyrus* Paperna, 1960 and *Scutogyrus* Pariselle & Euzet, 1995: *C. sclerosus* Paperna & Thurston, 1969, *C. halli* (Price & Kirk, 1967) Paperna, 1979, *C. tilapiae* Paperna, 1960, *C. thurstonae* Ergens, 1981, *C. cirratus* Paperna, 1964, *C. mbirizei* Muterezi Bukinga, Vanhove, Van Steenberge & Pariselle, 2012, *S. longicornis* (Paperna & Thurston, 1969) Pariselle & Euzet, 1995 and *S. gravivaginus* (Paperna & Thurston, 1969) Pariselle & Euzet, 1995 (Fig. 2).

**Fig. 2.**
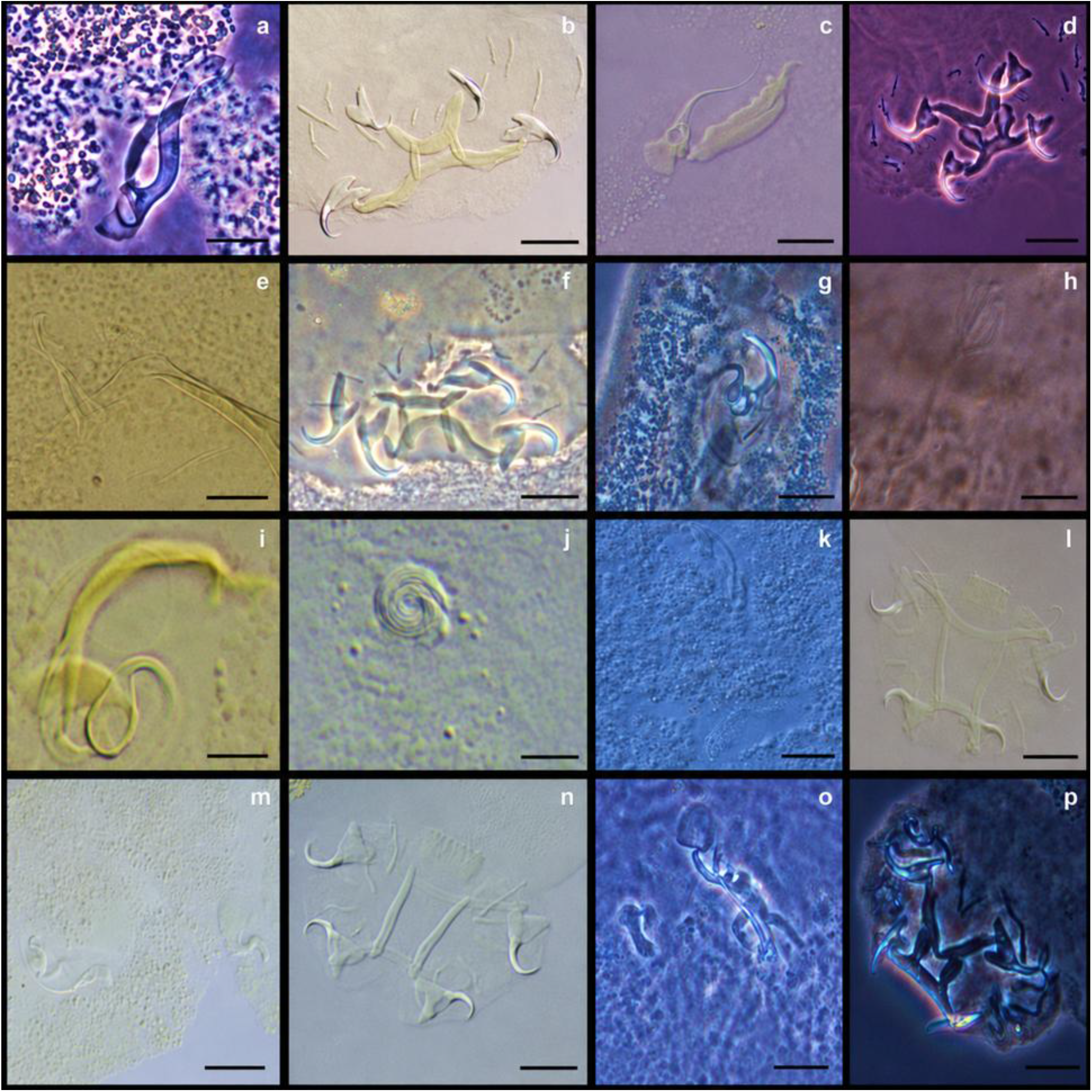
Photomicrographs of the copulatory and attachment organs of *C. halli* (a,b), *C. sclerosus* (c,d), *C. tilapiae* (e,f), *C. cirratus* (g,h), *C. mbirizei* (i,j), *S. longicornis* (k,l), *S. gravivaginus* (m,n) *and C. thurstonae* (o,p). Scale bars = 50 µm

Digenean metacercariae totalled 198 specimens in 27 fish; detailed species-level identification is reported in Shigoley et al. (2025). A total of 159 copepods were recovered from 34 fish. Molecular results supported assignment to *Lamproglena* with closest affinity to *L. monodi* (Supplementary Text S1; Figs. S1–S2). Sixteen acanthocephalans were recorded in ten fish and were identified morphologically as *Acanthogyrus* cf. *tilapiae* (Fig. 3). Molecular amplification did not yield usable products for acanthocephalans. Two nematodes were recorded and assigned to Anisakidae.

**Fig. 3.**
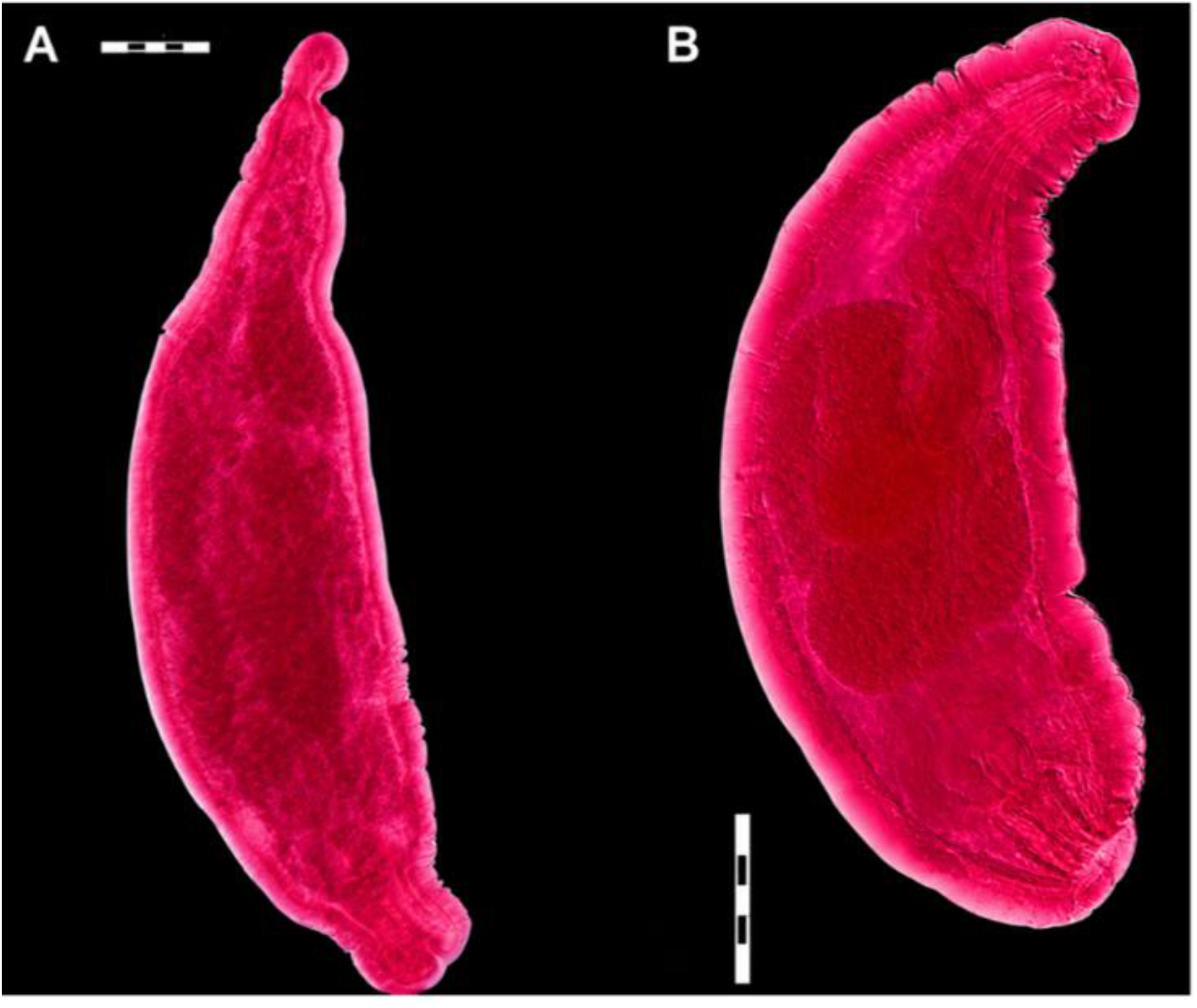
Light micrographs of whole-mounted acanthocephalans recovered from *Oreochromis niloticus*. (A) Female specimen with an elongated body; (B) male specimen with a broader, curved body. Scale bars: A = 200 µm; B = 500 µm

### 3.3. Exploratory parasite diversity across farm categories

Hill diversity was higher in the larger-scale category at all three orders: q = 0, 6.0 versus 1.0; q = 1, 3.20 versus 1.0; and q = 2, 2.62 versus 1.0 (Fig. 4). Wilcoxon rank-sum tests gave W = 60.5, p = 0.006 for q = 0; W = 61.0, p = 0.006 for q = 1; and W = 60.0, p = 0.008 for q = 2. Excluding F04 reduced the number of larger-scale farms from four to three, but the same pattern remained (q = 0, W = 44.5, p = 0.021; q = 1, W = 45.0, p = 0.018; q = 2, W = 44.0, p = 0.024). These comparisons describe the sampled operational categories and do not separate pond-number category from county or other farm characteristics.

**Fig. 4.**
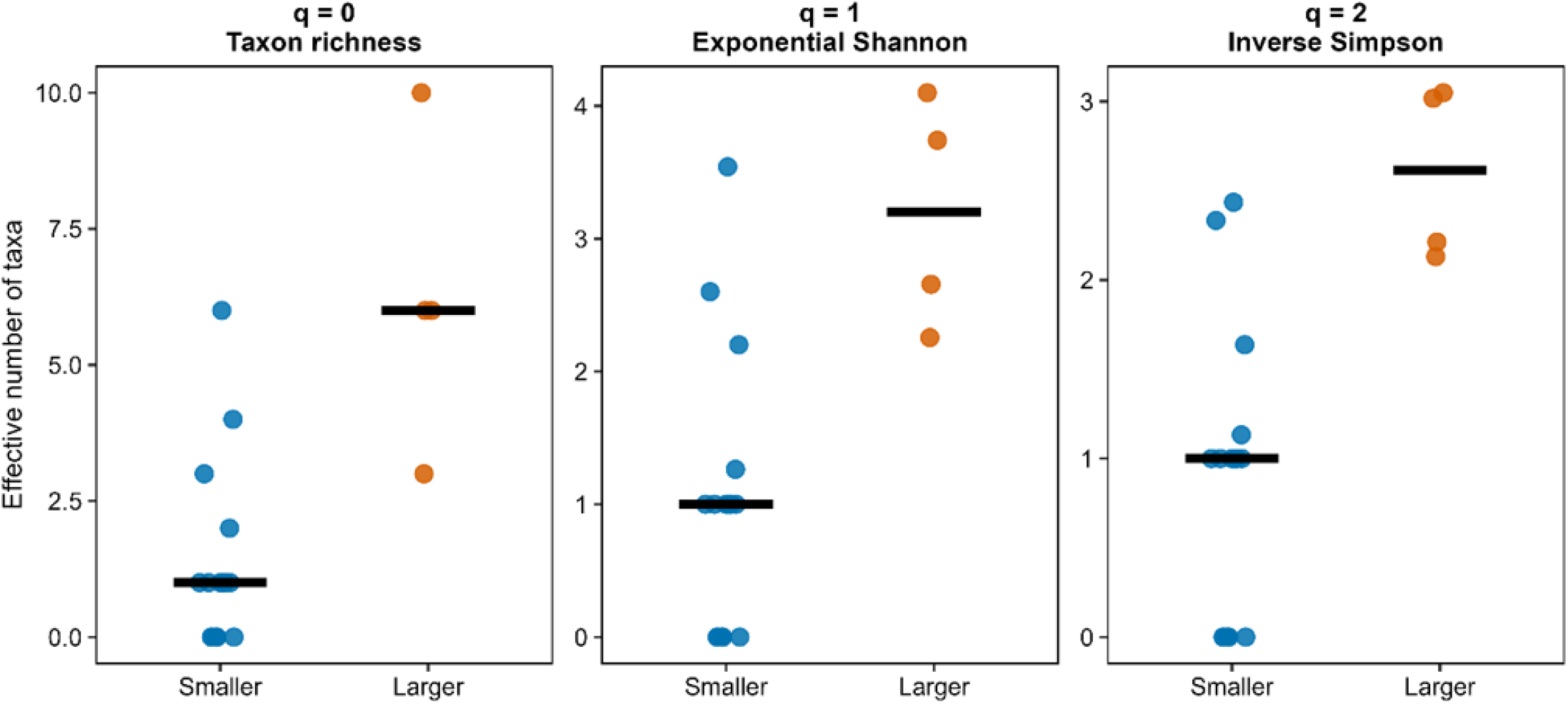
Farm-level parasite diversity expressed as Hill numbers by operational farm category. q = 0 represents taxon richness, q = 1 exponential Shannon diversity and q = 2 inverse Simpson diversity. Points represent individual farms and black horizontal bars indicate group medians. Parasite-negative farms were retained with diversity values of zero

Rarefaction gave a similar richness pattern after standardising the number of fish examined. At a common sampling effort of 72 fish, estimated richness was 12.0 taxonomic units in the larger-scale category and 6.0 in the smaller-scale category (Supplementary Fig. S3). After F04 was excluded, the categories could be compared at 22 fish; estimated richness was 10.0 and 5.14 taxonomic units, respectively.

### 3.4. Exploratory parasite community composition and water-quality context

Among the 14 parasite-positive farms, parasite-community composition differed between the sampled farm categories in the full analysis (PERMANOVA: pseudo-F = 3.31, R^2^ = 0.216, p = 0.034). After F04 was excluded, the estimated difference remained similar but statistical support weakened (pseudo-F = 2.40, R^2^ = 0.179, p = 0.055). Excluding F11 did not change the difference (pseudo-F = 3.45, R^2^ = 0.239, p = 0.022). Multivariate dispersion did not differ between categories in the full analysis (PERMDISP: p = 0.770). The first two PCoA axes explained 46.9% and 17.5% of the variation, respectively (Fig. 5).

**Fig. 5.**
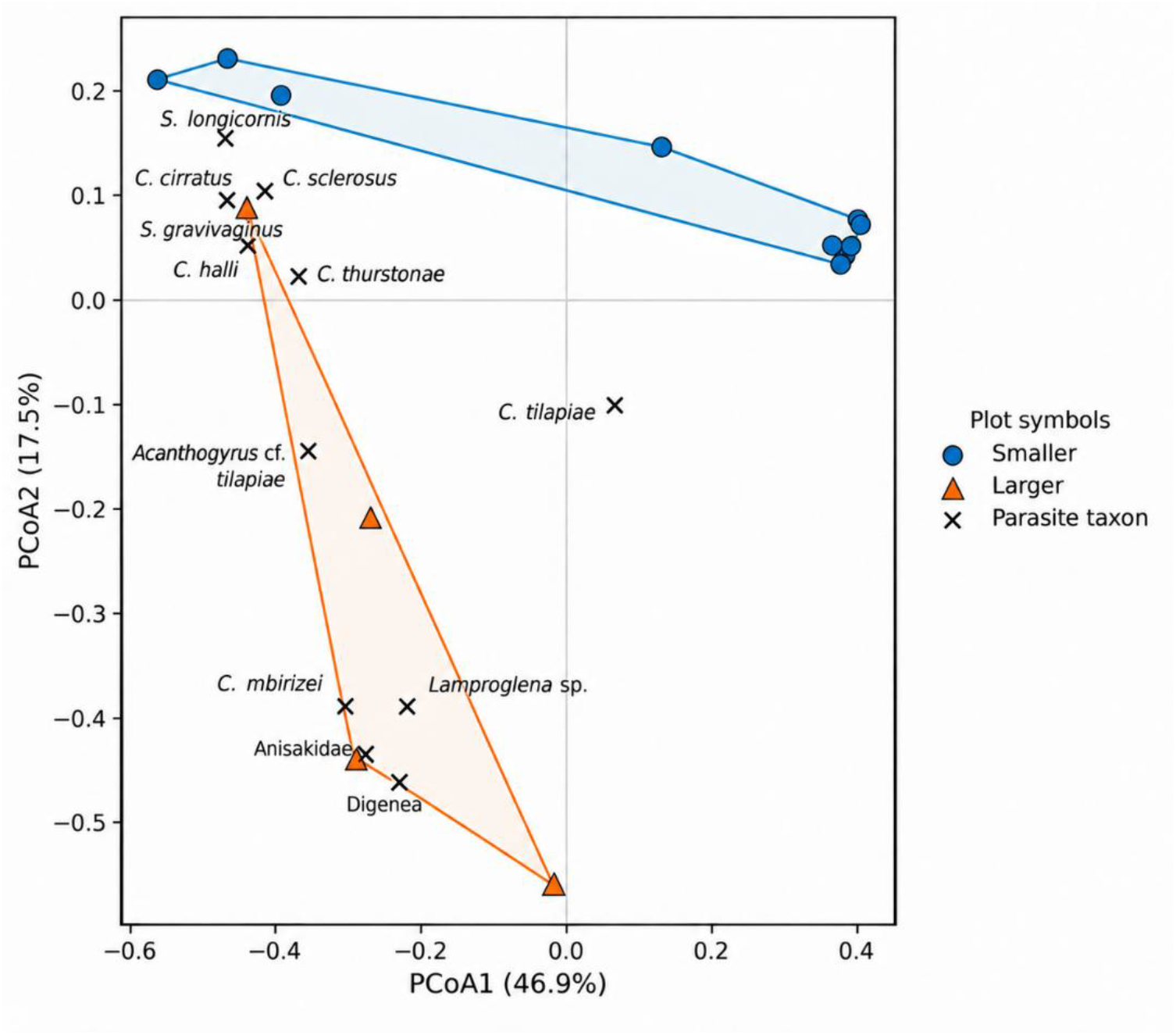
Principal coordinates analysis (PCoA) of farm-level parasite-community composition based on Bray–Curtis dissimilarities of square-root-transformed abundances of 12 parasite taxonomic units. Blue circles and orange triangles represent parasite-positive farms in the smaller- and larger-scale operational farm categories, respectively. Coloured convex hulls connect the outermost observed farms within each category, and black crosses indicate abundance-weighted taxon positions.

None of the four single-time-point water-quality measurements showed a clear association with farm-level parasite abundance after false-discovery-rate adjustment (Spearman |ρ| ≤ 0.144; FDR-adjusted p ≥ 0.999). Associations with the three Hill diversity measures were also unsupported (|ρ| ≤ 0.242; FDR-adjusted p ≥ 0.664). In the community analyses, individual water-quality variables explained 2.6–12.5% of the variation in parasite-community composition, but none was supported after FDR adjustment (PERMANOVA, adjusted p = 0.359–0.931). Sensitivity analyses excluding the influential farms gave the same overall conclusion. These measurements describe conditions at sampling and not seasonal or longer-term environmental exposure.

### 3.5. Gill lesions and parasite burden

Gill tissue from 137 of 157 fish was suitable for histological assessment. Epithelial hyperplasia was recorded in 124 fish, epithelial hypertrophy in 53, vascular congestion in 47, inflammation in 34 and lamellar fusion in 31 (Fig. 6). The cumulative number of lesion types ranged from 0 to 5, with a median of 2. Sixty-eight fish had no detected gill parasites, 17 had copepods only, 37 had monopisthocotylans only and 15 had both groups. In the primary ordinal model, increasing monopisthocotylan burden was associated with higher cumulative lesion-type categories (odds ratio = 1.86, 95% CI 1.43–2.42, p < 0.001). Copepod burden was not associated with the cumulative lesion-type count after adjustment for the other model terms (odds ratio = 1.16, 95% CI 0.72–1.88, p = 0.544), and neither host sex, farm category nor fish length was associated with the outcome. There was no evidence that the proportional-odds assumption was violated (nominal test, all p ≥ 0.100).

**Fig. 6.**
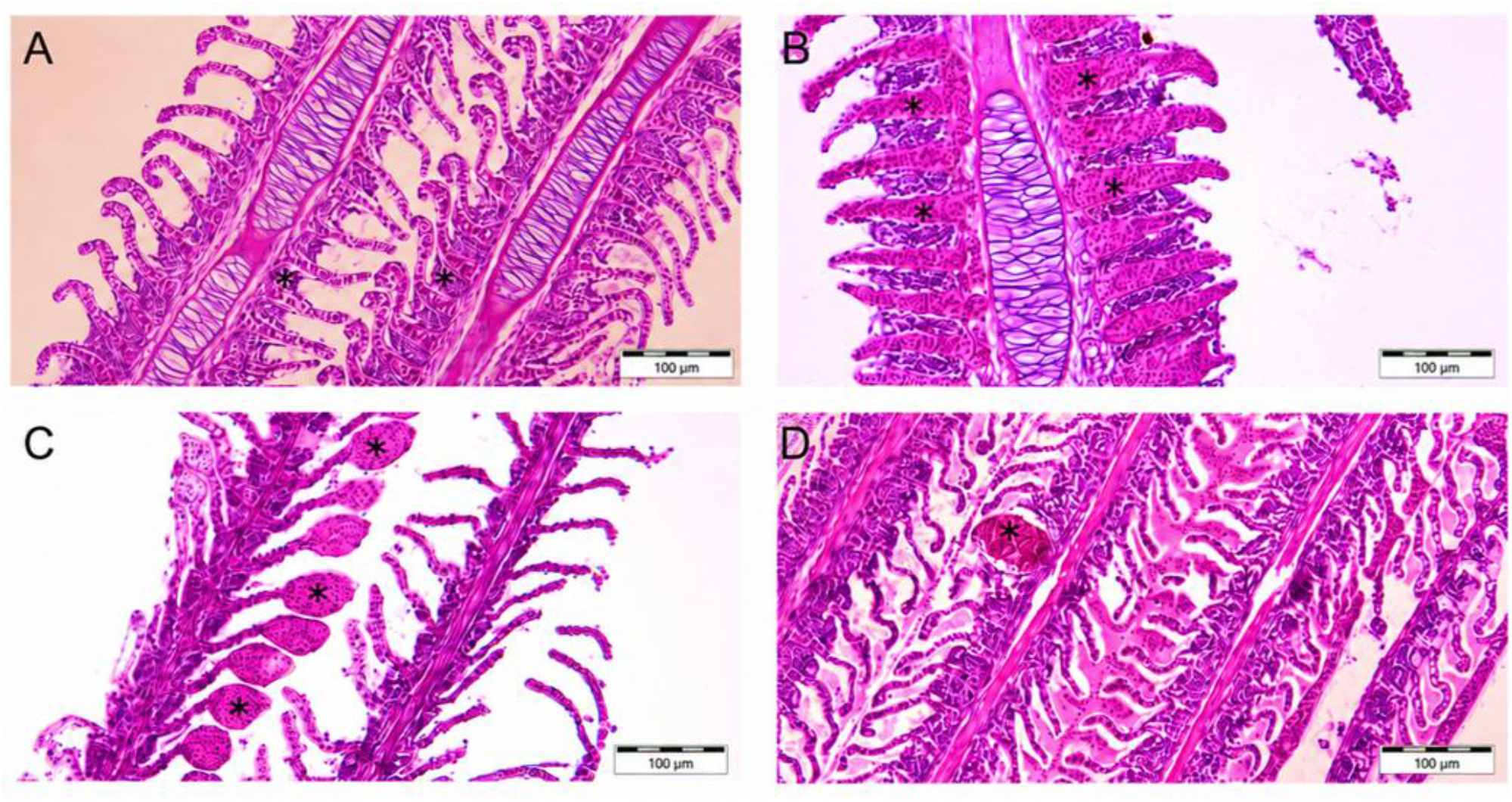
Representative photomicrographs showing: (A) epithelial hyperplasia, characterised by thickening of the secondary lamellar epithelium; (B) lamellar fusion, showing partial merging of adjacent secondary lamellae; (C, D) vascular congestion, represented by telangiectatic and aneurysmal dilation of lamellar blood spaces. Asterisks and arrows indicate the respective lesions. Scale bars: = 100 µm

When farm identity was included as a fixed effect, the monopisthocotylan estimate remained positive, but the 95% confidence interval included no association (odds ratio = 1.57, 95% CI 0.98–2.51, p = 0.059). Copepod burden remained unsupported. A secondary linear model with farm-clustered standard errors showed the same positive direction for monopisthocotylan burden (β = 0.377, p < 0.001). In lesion-specific analyses, monopisthocotylan burden was associated with epithelial hypertrophy, lamellar fusion and inflammation after correction for multiple testing; full results are given in Supplementary Table S2. These analyses indicate an association between monopisthocotylan burden and the number of gill lesion types but do not establish that the parasites caused each lesion.

### 3.6. Stakeholder recognition of fish-health problems and management responses

Among farmers, 27/36 (75%) reported having observed signs such as slow swimming, external lesions or discoloured gills. At the same time, 19/36 (53%) reported that they had not encountered disease and 4/36 (11%) were unsure, including some respondents who had experienced fish mortality. Thus, reporting abnormal signs and reporting a disease event were not equivalent in the questionnaire responses. Parasites were seldom named as a possible cause. Farmers described relying on fish behaviour, visible lesions and water colour to judge health and water conditions, and responses to suspected disease were often based on experience or trial and error rather than a confirmed diagnosis (Supplementary Table S1).

Farmers and fisheries or extension officers described difficulties accessing diagnostic support and reported little routine laboratory confirmation of fish-health problems. Participants also described gaps in fish-health training and feedback after diagnostic or research activities and expressed interest in further practical training. Representative statements are provided in Supplementary Table S3.

### 3.7. Integration of biological and stakeholder findings

The biological and stakeholder findings identified a diagnostic gap at study level. Targeted examination detected parasites and gill alterations, whereas farm-level assessment relied mainly on non-specific signs and rarely identified parasites as a specific cause. The exploratory pond-number comparisons and single-time-point water measurements did not identify the processes underlying parasite distribution. The combined findings are summarised in Table 3.

| Biological finding | Stakeholder observation | Interpretation |
|---|---|---|
| Monopisthocotylans occurred in 14/20 farms (10 smaller-scale and all four larger-scale farms), whereas copepods, digeneans, acanthocephalans and nematodes occurred only in the larger-scale farms | Parasites were rarely identified as a possible cause of fish-health problems | Parasite infections were detected through targeted examination but not consistently recognised at farm level |
| A higher monopisthocotylan burden was associated with a greater number of gill lesion types | Farmers reported discoloured gills, lesions, slow swimming and reduced feeding | Visible signs can indicate poor fish condition, but observation alone cannot identify the underlying cause |
| Single-time-point water measurements showed no clear association with parasite burden, diversity or community composition | Water conditions were commonly judged from colour and fish behaviour | Both approaches provide limited information on longer-term water conditions; repeated measurements would be needed to assess environmental exposure |
| Parasite detection and lesion assessment required microscopy and histology, with molecular analysis supporting selected identifications | Farmers and officers reported limited access to diagnostic services, training, and feedback | Routine farm observations were not consistently linked to laboratory confirmation or specific diagnosis |

## 4. Discussion

The main fish-health finding of this study was that higher monopisthocotylan burden was associated with a greater number of gill lesion types, while farmers and fisheries or extension officers rarely identified parasites as a possible cause of poor fish health. Together, these findings point to a gap between laboratory detection and farm-level diagnosis. Parasite diversity and community composition also differed between operational pond-number categories, but these farm-level patterns were exploratory because only four farms were in the larger category, category overlapped with county and individual farms strongly influenced the results. Single-time-point water measurements showed no clear associations with parasite burden, diversity or community composition.

Parasite diversity was higher in the larger pond-number category at q = 0, q = 1 and q = 2, and the pattern remained after F04 was excluded. Rarefaction similarly retained a richness difference after standardising the number of fish examined. Community composition differed between categories in the full PERMANOVA, but support weakened after F04 was excluded. These results show a pattern among the sampled farms rather than an independent effect of production scale: the category was based only on pond number, three of the four larger farms were in Kiambu, and it did not capture stocking density, water source, fish introductions, biosecurity or other management characteristics that may affect parasite transmission.

Water conditions measured at sampling varied among farms. Temperature ranged from 22.1 to 28.0 °C, pH from 6.7 to 10.2, TDS from 15.5 to 96.0 mg L^−1^ and conductivity from 34.0 to 651 µS cm^−1^, with particularly wide between-farm variation in conductivity. For Nile tilapia, optimal rearing temperatures of approximately 27–32 °C and pH values around 7–8 have been reported (Mengistu et al., 2020). Temperatures were below this optimum at 14 of the 20 farms, and pH was above 8 at 16 farms and below 7 at two. Similar variation has been reported from fish ponds in Kirinyaga County, where temperatures ranged from 20.5 to 31.7 °C and pH from 6.27 to 9.2 (Wanja et al., 2020). Despite this variation, none of the single-time-point measurements showed a clear association with parasite abundance, diversity or community composition after correction for multiple testing. These measurements characterise conditions at sampling only and cannot capture seasonal variation or environmental conditions during earlier stages of parasite transmission. Repeated measurements across seasons and production cycles would therefore be needed to examine environmental exposure more directly.

The gill findings provide the strongest biological evidence of a fish-health association in this study. Monopisthocotylan burden was positively associated with the number of lesion types, and secondary analyses linked monopisthocotylan burden with epithelial hypertrophy, lamellar fusion and inflammation. These alterations are relevant changes in gill structure but are not specific to parasites and can arise from other stressors (Bernet et al., 1999; Steckert et al., 2018). When farm identity was included as a fixed effect, the estimated monopisthocotylan association remained positive, but the 95% confidence interval included no association, indicating that between-farm differences may contribute to the observed relationship. The findings therefore support an association between heavier monopisthocotylan infection and a greater number of gill alterations in the sampled fish, not proof that parasites caused each lesion.

The stakeholder findings show that recognition of poor fish condition did not necessarily translate into recognition of a disease event or identification of its cause. Farmers frequently reported slow swimming, lesions, discoloured gills and mortality, yet many also reported that they had not encountered disease. Parasites were seldom identified as a possible explanation. Visible abnormalities, mortality and “disease” were therefore not necessarily equivalent categories in participants’ accounts. Similar gaps in access to aquatic animal-health expertise, diagnostic services and biosecurity support have been reported elsewhere in African aquaculture (Brugere et al., 2017; Kyule-Muendo et al., 2022).

Management responses reflected this diagnostic uncertainty. Farmers described relying on fish behaviour, visible lesions and water colour to assess fish and pond condition, while treatment decisions could proceed through trial and error without laboratory confirmation. Participants also reported limited access to diagnostic feedback and expressed interest in further practical fish-health training. The veterinary officer interviewed noted that fish were not treated in the same way as other livestock and that fish health had received only brief coverage in their veterinary curriculum. This account suggests that gaps in fish-health training may also affect veterinary professionals. The main challenge was therefore not simply recognising that fish were unwell, but moving from observation of non-specific signs to identification of the underlying cause and an informed response. Strengthening sample-referral pathways, diagnostic feedback and practical training for farmers, extension officers and veterinary professionals could help address these gaps.

The biological study design limits causal interpretation. Only 20 farms were sampled, and four were in the larger pond-number category. Farm category overlapped with county, sampling effort was unequal, particularly at F04, and F11 had an exceptionally high monopisthocotylan burden. Water quality was measured once per farm, and gill alterations were recorded as present or absent and were not parasite-specific. These limitations restrict inference about independent effects of pond-number category, longer-term environmental exposure and parasite causation.

The findings support practical improvements in fish-health assessment. When farms experience persistent behavioural changes, gill abnormalities or unexplained losses, affected fish could be referred for histopathology or other parasite screening rather than relying on visual signs alone. Referral pathways should include clear procedures for sample collection and feedback of diagnostic results to farmers and extension personnel. Future studies should sample more farms within each county, collect repeated water-quality and management data and examine parasite-specific tissue effects while accounting for co-infections.

## 5. Conclusion

Higher monopisthocotylan burden was associated with a greater number of gill lesion types in farmed Nile tilapia from central Kenya, although the farm-fixed-effect sensitivity analysis was less conclusive. Farmers and fisheries or extension officers recognised signs of poor fish health but seldom identified parasites as a possible cause, and routine diagnostic confirmation was uncommon. Parasite diversity and community composition differed between operational pond-number categories, but these exploratory comparisons were limited by the small and geographically concentrated larger category. Single-time-point water-quality measurements showed no clear associations with parasite patterns. Strengthening parasite screening, sample-referral procedures and diagnostic feedback could better connect farm observations with laboratory evidence.

## CReDiT authorship contribution statement

**Miriam Isoyi Shigoley**: Conceptualization, Methodology, Investigation, Formal analysis, Visualisation, Writing – original draft, Writing – review & editing. **Nicolas Antoine-Moussiaux**: Project supervision, Methodology, Writing – review & editing. **Thierry Jauniaux**: Project supervision, Investigation, Writing – review & editing. **Thomas-Julian Irabor**: Methodology, Formal analysis, Writing – review & editing. **Gyrhaiss Kapepula Kasembele**: Formal analysis, Writing – review & editing. **Daniel Mungai Ndegwa:** Investigation, Writing – review & editing. **Bram J. Vanschoenwinkel**: Formal analysis, Writing – review & editing. **Maarten P. M. Vanhove**: Project supervision, Investigation, Writing – review & editing.

## Declaration of Competing Interest

The authors declare they have no competing interests.

## Acknowledgements

We would like to thank all the farmers, fisheries and extension officers who participated in the survey. Edward Njagi, Gilbert Kosgei, Gathua Joseph and Justus Ochong’ (Ichthyology section, National Museums of Kenya) are thanked for supplying field equipment and the storage of tissue samples. Tom Artois and François Mélard are thanked for their contributions to developing this research project.

## Funding

This research was funded by the BOF22DOCLI04 - Special Research Fund of Hasselt and Liège University and the International and Intersectoral mobility funding - 2024, Hasselt University. Part of the research leading to the results presented in this publication was carried out with infrastructure funded by the European Marine Biological Research Centre (EMBRC) Belgium, Research Foundation – Flanders (FWO-Vlaanderen) (GOH3817N).

## Notes

### Competing Interest Statement

The authors have declared no competing interest.

