## Supplementary Materials for "Hidden infections: Linking parasite ecology, fish health and stakeholder perceptions in Nile tilapia aquaculture in central Kenya"

Table S1. Summary of recurring stakeholder observations on fish-health recognition and management during the main field study

| Theme | Observation | Evidence in the field study | Interpretation |
| --- | --- | --- | --- |
| Recognition of poor health | Reliance on visible and behavioural signs | 27/36 farmers (75%) reported signs such as slow swimming, lesions or discoloured gills | Farmers could recognise that fish were unwell, but signs were non-specific |
| Perception of disease | Disease was not always recognised despite health events | 19/36 farmers (53%) reported no disease and 4/36 (11%) were unsure, including some respondents who had experienced mortality | Health problems were not consistently described as disease events |
| Parasite awareness | Parasites were rarely named as a cause | Parasites were seldom mentioned without prompting | Specific parasite recognition was limited |
| Diagnostic approach | Visual inspection predominated | Fish behaviour, lesions and water colour were commonly used to judge health | Routine laboratory confirmation was uncommon |
| Disease response | Treatment was often empirical | Farmers described isolating fish or trying treatments without a confirmed diagnosis | Management decisions were often made before the cause was known |
| Water assessment | Water colour and fish behaviour were common indicators | Instrument-based routine monitoring was not consistently reported | Visual assessment could not describe longer-term water chemistry |

Table S2. Secondary binary logistic-regression results for individual gill lesions. Odds ratios refer to a one-unit increase in ln(parasite abundance + 1). Models also included host sex, operational farm category and standardised fish length. False-discovery-rate adjustment was applied separately across the five lesion models for each parasite predictor

| Lesion | Parasite predictor | Odds ratio | 95% CI | p-value | FDR-adjusted p |
| --- | --- | --- | --- | --- | --- |
| Epithelial hypertrophy | Monopisthocotylans | 2.04 | 1.40–2.96 | <0.001 | <0.001 |
| Epithelial hyperplasia | Monopisthocotylans | 1.04 | 0.66–1.66 | 0.858 | 0.976 |
| Lamellar fusion | Monopisthocotylans | 1.58 | 1.16–2.16 | 0.004 | 0.006 |
| Inflammation | Monopisthocotylans | 2.32 | 1.60–3.35 | <0.001 | <0.001 |
| Vascular congestion | Monopisthocotylans | 1.00 | 0.74–1.33 | 0.976 | 0.976 |

| | | | | | |
|---|---|---|---|---|---|
| Epithelial hypertrophy | Copepods | 2.40 | 1.13–5.09 | 0.023 | 0.090 |
| Epithelial hyperplasia | Copepods | Not estimable | — | — | — |
| Lamellar fusion | Copepods | 0.62 | 0.29–1.32 | 0.216 | 0.288 |
| Inflammation | Copepods | 0.99 | 0.48–2.02 | 0.972 | 0.972 |
| Vascular congestion | Copepods | 0.64 | 0.35–1.17 | 0.149 | 0.288 |

For copepods and epithelial hyperplasia, the standard logistic model could not estimate a finite effect because all 32 copepod-positive fish had hyperplasia. This result is therefore not interpreted.

Table S1. Representative stakeholder statements illustrating fish-health perceptions and management practices.

| **Theme** | **Representative statement** | **Interpretation** |
|---|---|---|
| Disease perception | A sick fish dies | Disease described mainly in terms of outcome rather than specific cause |
| Knowledge of diseases and parasites | I do not know the exact names of diseases or parasites | Limited ability to name specific fish-health problems |
| Possible source of health problems | I think the source of my fingerlings could be a problem | Possible recognition that fish source may influence health risk |
| Seasonal observations | When it gets colder, I see fish dying one by one | Seasonal observations were used to interpret mortality |
| Water assessment | If the water is too green, it is time to change it | Water conditions were judged visually |
| Treatment decisions | I try one treatment, then another if it fails | Treatment could proceed without a confirmed diagnosis |
| Technical support | Officers cannot tell us what caused the problem | Perceived limits in access to specific diagnostic advice |
| Information flow | We never get feedback | Reported lack of feedback after investigations or research |
| Training | I would like to join training on fish health | Interest in further practical fish-health training |

**Supplementary Methods: Copepod DNA extraction and PCR conditions**

DNA was extracted from copepod egg strings using a protocol adapted from Kmentová et al. (2021). Samples stored in 99% ethanol were dried for 30 min, then incubated overnight at 55 °C in 195 µL TNES buffer (400 mM NaCl, 20 mM EDTA, 50 mM Tris pH 8, 0.5% SDS) and 5 µL proteinase K (20 mg/mL).

After addition of 2 µL yeast tRNA (10 mg/mL), 65 µL 5 M NaCl and 290 µL 96% ethanol were added. Samples were cooled for 60 min at −20 °C, centrifuged for 15 min at 18,000 rcf, washed twice with chilled 70% ethanol and eluted in 30 µL 0.1× TE buffer. DNA was resuspended overnight at 4 °C and stored at −20 °C.

For 18S and 28S rDNA, 25-µL PCR reactions contained 2 µL DNA extract, 12.50 µL MangoMix™, 0.50 µL $MgCl_2$ (1 mM), 1.25 µL each of the forward and reverse primers (0.5 µM) and 7.50 µL dd$H_2O$. Cycling consisted of 2 min at 94 °C; 39 cycles of 1 min at 94 °C, 1 min at 50 °C and 1 min 30 s at 72 °C; followed by 7 min at 72 °C. COI amplification used LCO1490/HCO2198 with 5 min at 95 °C; 37 cycles of 30 s at 95 °C, 30 s at 47 °C and 1 min at 72 °C; followed by 7 min at 72 °C.

**Supplementary Text S1: Molecular identification of copepods**

Molecular identification was performed on representative copepod specimens using 28S rDNA and COI mtDNA. For 28S, 24 selected specimens were processed and 23 produced good-quality sequences. Most retained sequences were identical across the analysed fragment; three representative sequences (COP3_present study, COP16_present study and COP19_present study) were therefore retained for the phylogenetic comparison. Pairwise divergence among these three sequences ranged from 0.0000 to 0.0014 substitutions per site. Their closest matches among the reference sequences were *Lamproglena monodi*, supporting assignment of the specimens to *Lamproglena* with closest affinity to *L. monodi*. The corresponding maximum-likelihood tree is shown in Supplementary Fig. S1.

Four COI sequences were retained for comparison. Pairwise divergence among the present-study COI sequences ranged from 0.0327 to 0.0892 substitutions per site. The smallest observed distance to a reference sequence was 0.0015 between one present-study sequence and *L. monodi* OR058788.1. In the maximum-likelihood tree, one present-study sequence clustered with *L. monodi*, while two other sequences formed a sister pair (Supplementary Fig. S2). Because support differed between markers and not all present-study COI sequences grouped directly with the reference, the copepods are reported in the manuscript as *Lamproglena* sp. with closest molecular affinity to *L. monodi*.

**Supplementary Interview Guide S1: Semi-structured interview guide used for stakeholder interviews**

*The interview aims to gather valuable insights on what people think about Nile tilapia health. Your responses will remain confidential, and no personal information will be disclosed.*

1. Stakeholder role: What role (s) do you play as a stakeholder in Nile tilapia production?

2. Production challenges: What problems hinder Nile tilapia production?

3. Prioritising challenges: From the problems you have mentioned, which ones do you prioritise so far?

4. Disease experience: Have you ever experienced a fish getting sick? If yes, what do you think caused it? How often does it happen?

5. Early warning indicators of disease: Are there any early signs you've noticed that show fish might be getting sick? Can you describe them?

6. Preparation for disease outbreak: Do you do anything in advance to protect your fish from getting sick? If so, what do you do?

7. Response to disease outbreak: In the event of a disease outbreak, can you describe how you detect, respond to and recover from a disease crisis?

8. Information needs: What information or knowledge about fish health and disease is missing?

9. Collaboration for disease management: What sectors or stakeholders do you believe need to collaborate for disease management in Nile tilapia?

10. Recommendation for improving Nile tilapia health: What advice or suggestions do you have for improving the health and production of Nile tilapia?
