## Supplementary figures and images for "Hidden infections: Linking parasite ecology, fish health and stakeholder perceptions in Nile tilapia aquaculture in central Kenya"

### Supplementary Figure S1

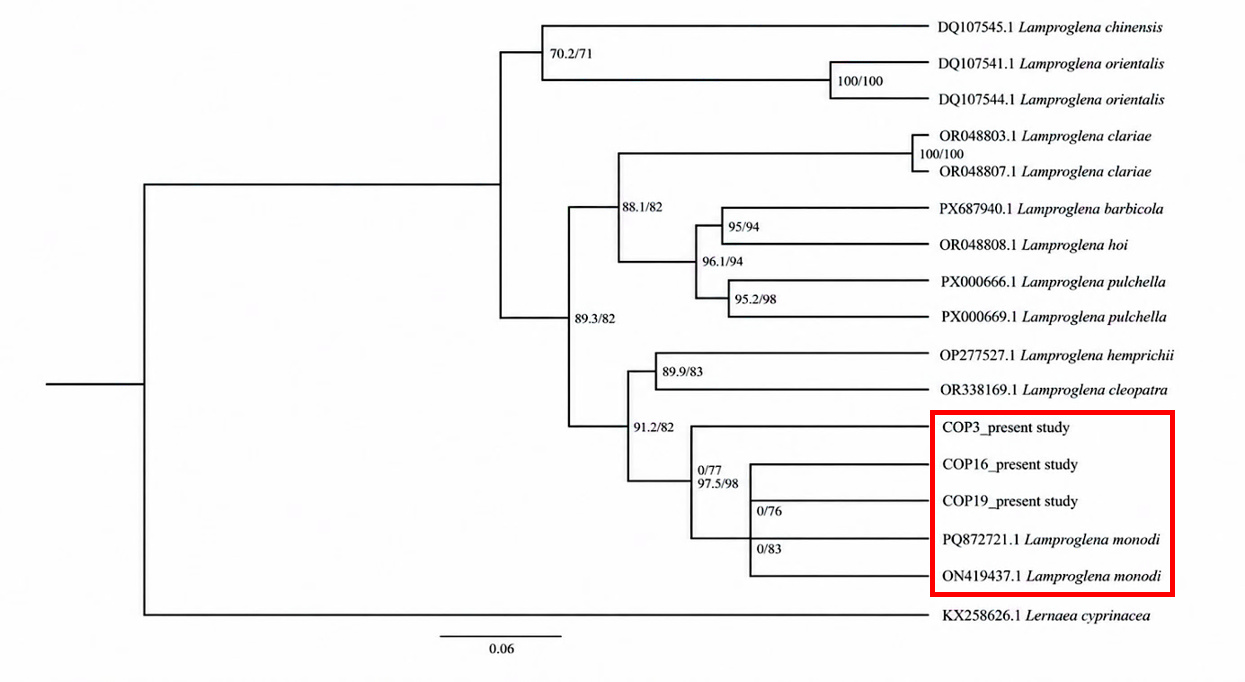

### Supplementary Figure S2

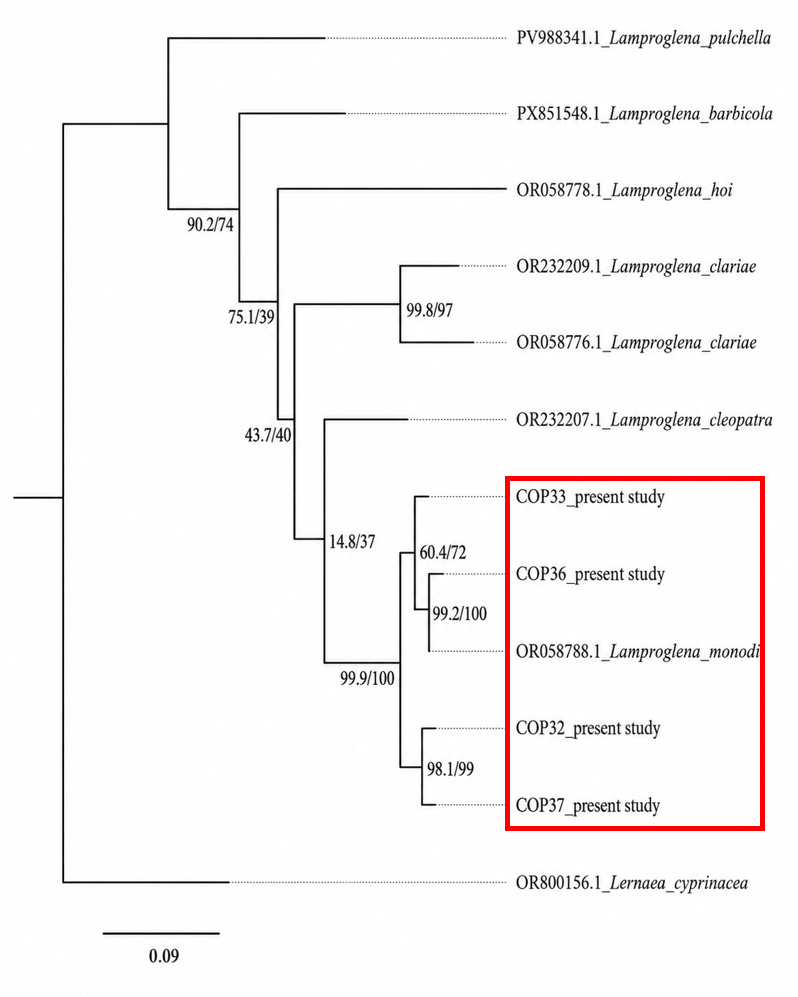

### Supplementary Figure S3

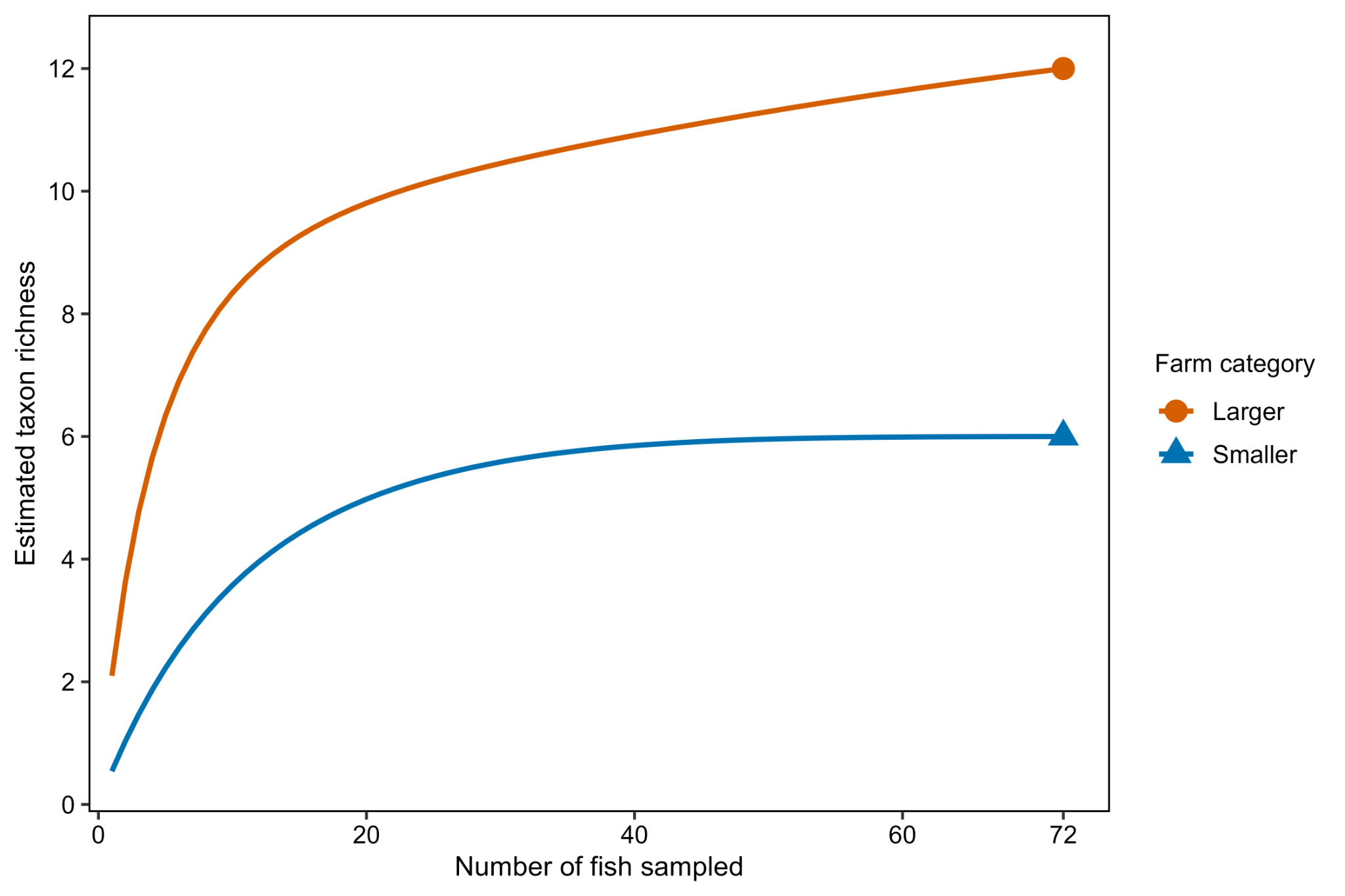
